# Human sialomucin CD164 is an essential entry factor for lymphocytic choriomeningitis virus

**DOI:** 10.1101/2022.01.24.477570

**Authors:** Jamin Liu, Kristeene A. Knopp, Elze Rackaityte, Chung Yu Wang, Matthew T. Laurie, Sara Sunshine, Andreas S Puschnik, Joseph L DeRisi

**Affiliations:** Department of Biochemistry and Biophysics, University of California, San Francisco, San Francisco, California, USA; University of California, Berkeley-University of California, San Francisco Graduate Program in Bioengineering, University of California, San Francisco, San Francisco, California, USA; Chan Zuckerberg Biohub, San Francisco, California, USA

**Author notes:** Address correspondence to Joseph L. DeRisi.

## Abstract

Lymphocytic choriomeningitis virus (LCMV) is a well-studied mammarenavirus that can be fatal in congenital infections. However, our understanding of LCMV and its interactions with human host factors remain incomplete. Here, host determinants affecting LCMV infection were investigated through a genome-wide CRISPR knockout screen in A549 cells, a human lung adenocarcinoma line. We identified and validated a variety of novel host factors that play a functional role in LCMV infection. Among these, knockout of the sialomucin CD164, a heavily glycosylated transmembrane protein, was found to ablate infection with multiple LCMV strains but not other hemorrhagic mammarenaviruses, in several cell types. Further characterization revealed a dependency of LCMV entry on the cysteine-rich domain of CD164, including a N-linked glycosylation site at residue 104 in that region. Given the documented role of LCMV with respect to transplacental human infections, CD164 expression was investigated in human placental tissue and placental cell lines. CD164 was found to be highly expressed in the cytotrophoblast cells, an initial contact site for pathogens within the placenta, and LCMV infection in placental cells was effectively blocked using a monoclonal antibody specific to the cysteine-rich domain of CD164. Together, this study identifies novel factors associated with LCMV infection of human tissues, and highlights the importance of CD164, a sialomucin that has previously not been associated with viral infection.

## Introduction

The Arenaviridae family is classified into four genera: *Antennavirus* which were discovered in actinopterygian fish; *Reptarenavirus* and *Hartmanivirus* which infect boid snakes; and *Mammarenavirus* whose hosts are predominantly rodents(1–4). Mammarenavirus can be further divided into two major virus subgroups based on antigenic properties: Old World (OW), which are mainly indigenous to Africa, and New World (NW), which are indigenous to the Americas(5). Several viruses from this genus can also infect humans, leading to severe or fatal disease. One such pathogenic mammarenavirus is lymphocytic choriomeningitis virus (LCMV). Considered to be the prototypic arenavirus, LCMV is an OW virus found on all populated continents due to the ubiquitous distribution of its natural host, the house mouse (*Mus musculus*)(6). The prevalence, however, among humans as measured through serological presence of LCMV antibodies widely varies (from 4% to 13%), making it challenging to estimate disease burden and infection risk(7). In addition to contact with infected rodents, humans can also become infected with LCMV through solid organ transplant or by vertical transmission. In the former case, LCMV infection in immunosuppressed organ recipients is frequently fatal and the only available therapeutic is off-brand use of nucleoside analog ribavirin (8). As for the latter, transplacental infection leading to congenital LCMV are typically abortive or result in severe and often fatal fetopathy(9, 10).

Like all other mammarenaviruses, LCMV is a pleiomorphic enveloped virus with a bi-segmented, ambi-sense, negative-stranded RNA genome encoding four genes(11). The L segment (7.2 kb) encodes the viral RNA-dependent RNA polymerase (L), and a small RING finger protein (Z) that is functionally equivalent to the matrix protein found in many enveloped RNA viruses. The S segment (3.4 kb) encodes the viral nucleoprotein (NP) and the glycoprotein complex (GPC). The GPC is synthesized in the infected cell as a precursor polypeptide before being proteolytically processed into a stable signal peptide (SSP), and two noncovalently linked subunits GP1 and GP2 by the protease SKI-1/S1P(12). GP1 subunit associates with a cellular receptor while GP2 is a transmembrane protein that mediates the pH dependent fusion of viral and cellular membranes in the late-stage endosomes(13–15). All three subunits remain associated in a tripartite complex while expressed on the viral surface to facilitate viral attachment and entry(12, 16).

Dystroglycan (DAG1), a widely expressed cell adhesion molecule, is recognized as the main attachment factor for viral entry by LCMV, Lassa Virus (LASV) and several other NW mammarenaviruses(17, 18). DAG1 is expressed as a precursor polypeptide that is post-translationally cleaved into two noncovalently associated subunits, the peripheral membrane alpha subunit (αDG) and the transmembrane beta subunit (βDG)(19). Additionally, αDG undergoes complex O-glycosylation mediated by the glycotransferase like-acetylglucosaminyl-transferase (LARGE). Appropriate LARGE-dependent glycosylation is critical for interaction between αDG and mammarenavirus GP(20, 21).

LCMV cellular tropism, however, does not always correlate with the presence of fully glycosylated αDG and certain strains of LCMV are still found to efficiently bind and infect host cells in the complete absence of DAG1(22). Previous studies have shown that single amino acid substitutions such as S153F, Y155H, and L260F in the GP1 domain can alter the binding affinity to αDG and shift GP binding preference to alternative receptors(23). This allowed for further classification of LCMV strains into high- and low-αDG LCMV variants. Several secondary receptors have been proposed, including members of the Tyro3/Axl/Mer (TAM) family and heparan sulfate proteoglycans(24–26). Interestingly, in each case, residual viral infection is still observed in when tested in genetic knockouts, implying the presence of additional receptors able to mediate cell entry.

The cell entry process reaches completion for mammarenaviruses when viral and cell membrane fusion allows the viral RNP to be deposited into the cytoplasm. For OW mammarenaviruses Lassa virus and Lujo virus, this step requires GP2 to bind to late endosomal resident proteins LAMP1 and CD36, respectively, in a low pH environment(27, 28). Whether LCMV also requires such a receptor switch in the late endosome is currently unknown.

Although LCMV is considered the prototypic mammarenavius and is consistently used as a model to study the effect of viral persistence on host immunity, several aspects of its viral life cycle and cellular tropism remain incompletely understood. In this study, we explored the essential host requirements for LCMV infection by performing a genome-wide CRISPR Cas9 knockout (KO) screen using the GeCKOv2 guide library(29). Our results identify new host factors associated with LCMV infection, while also corroborating previously implicated factors. Among these results, we identify CD164 as an essential entry factor and possible therapeutic target for LCMV infection.

## Results

### CRISPR KO screens identify host factors for LCMV infection

LCMV is a virus with minimal cytopathic effect. To conduct a genome-wide pooled CRISPR KO screen to identify host factors important for LCMV infection, a recombinant tri-segmented LCMV reporter virus (rLCMV-mCherry) with one L segment and two S segments was constructed(30). We genetically encoded mCherry in place of the nucleoprotein (NP) on one S segment and in place of the glycoprotein complex (GPC) on the other S segment (**Figure S1A**). One-step growth curves demonstrated slower growth kinetics for rLCMV-mCherry compared to its parental strain, LCMV Armstrong 53b (Arm 53b), with final titers being comparable (**Figure S1B**). 24 hours post infection (hpi), the percentage of cells expressing mCherry (94.2% mCherry+) was equivalent to the percent expressing nucleoprotein (99.6% N protein+) suggesting minimal deleterious effects of this tri-segmented genome arrangement (**Figure S1C**).

As inhalation of aerosolized virus is a major transmission route, human adenocarcinoma lung epithelial cells (A549) were the chosen cell line for whole-genome CRISPR KO screening with the GeCKOv2 guide library. Following rLCMV-mCherry infection (Multiplicity of infection (MOI) 10) of the A549 CRISPR KO library cells, mCherry-negative cells were sorted 24 hours post infection (hpi) to select for single-guide RNAs (sgRNAs) targeting host factors necessary for successful LCMV infection (**Figure 1A**). The sgRNAs present in this virus-resistant and an unsorted control population were PCR amplified from the extracted genomic DNA and subsequently identified via next-generation sequencing. Using the MAGeCK algorithm, genes were ranked using robust rank aggregation to produce a significance score called the MAGeCK enrichment score(31). As expected, multiple sgRNAs targeting the same gene were among the top scoring guides, including those targeting previously described mammarenavirus host factors (**Figure 1B, Figure S2A, Table S1**). These include sialic acid metabolism genes (*ST3GAL4, SLC35A1*) and glycosylation related genes (Conserved Oligomeric Golgi (COG) complex members, *TMEM165*) which have been shown to be LASV host factors(32). Multiple heparan sulfate biosynthetic genes (*EXTL3, NDST1, PTAR1, SLC35B2*) described to be relevant for Lujo virus and DAG1-independent LCMV infections were also enriched(25, 28). The LCMV attachment factor *DAG1* was detected, albeit at a lower enrichment score. Additional host factors that were significantly enriched include those described for other viral infections, such as negative-stranded RNA virus vesicular stomatitis virus (VSV) (*ARFRP1, SYS1, YKT6*) and the human immunodeficiency virus (HIV) (*SRP14, DYRK1A, IL2RA*)(33–37).

**Figure 1.**
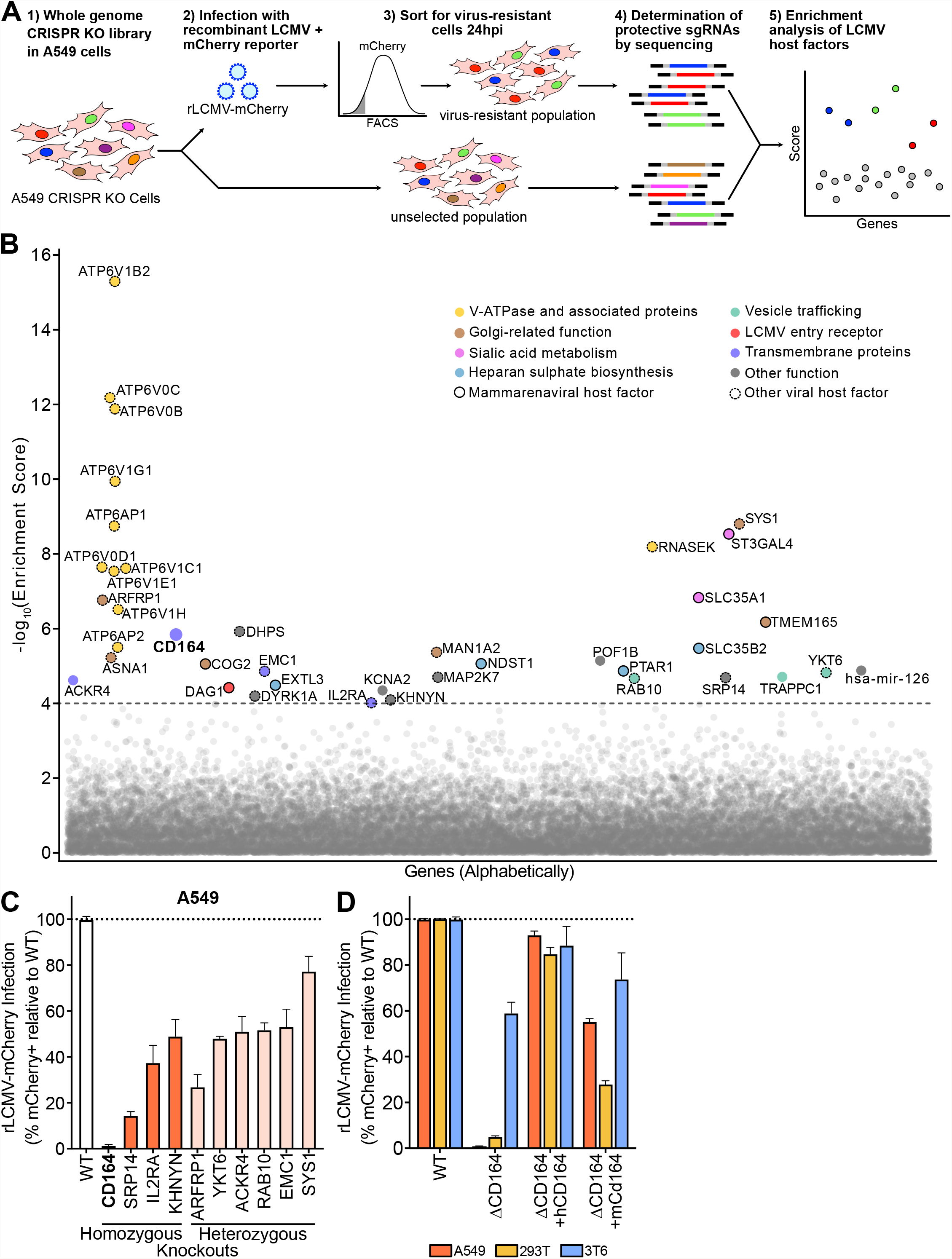
Genome-wide CRISPR loss-of-function screen in human cells identify host factors important for LCMV infection. (A) Schematic of CRISPR-based KO screen done in A549 lung epithelial cells for the identification of LCMV host factors. (B) Gene enrichment for CRISPR screen of rLCMV-mCherry infection. Enrichment scores were determined by MaGECK analysis and genes were colored by biological function. Dotted line indicates −log10(Enrichment Score) = 4. All genes and their enrichment scores can be found in **Table S1**. (C) Percentage of infected cells as determined by flow cytometry following infection of A549 homozygous knockouts (CD164, SPR14, IL2RA, KHNYN) or heterozygous knockouts (ARFRP1, YKT6, ACKR4, RAB10, EMC1, SYS1) with rLCMV-mCherry. Wildtype cells were used as normalization controls. Cells were infected at MOI 1 and harvested at 24 hpi. Error bars indicate standard error of three independent experiments. (D) Quantification of viral infection in WT, Δ*CD164*, Δ*CD164* complemented with human *CD164* (Δ*CD164* + *hCD164*), and Δ*CD164* complemented with mouse *Cd164* (Δ*CD164* + *mCd164*) in A549, 293T, and 3T6 cell type backgrounds. Cells were infected with rLCMV-mCherry at MOI 1 and harvested at 24 hpi. Error bars indicate standard error of three independent experiments.

Gene Ontology (GO) overrepresentation analysis of the top 300 hits from the screen using PANTHER(38) indicated an enrichment of genes associated with the signal recognition particle (*SRP14, SRP68, SRP19, SRP72*) and proton transmembrane transporter activity (*ATP6V1E1, ATP6V0D1, ATP6V1B2, ATP6V0C, ATP6V1A, ATP6V1G1, ATP6V0B, ATP12A, ATP6V1H, ATP6V1C1, CLCN4, ATP6V1F, ATP5S*) (**Table S2**). These same hits are also overrepresented in GO cellular components signal recognition particle and vacuolar proton-transporting vacuolar type ATPase (v-ATPase) complex, respectively.

Nearly every subunit of the v-ATPase (*ATP6V1B2, ATP6V0C, ATP6V0B, ATP6V1G1, ATP6AP1, ATP6V0D1, ATP6V1C1, ATP6V1E1, ATP6V1H, ATP6AP2*) was enriched in our screen. V-ATPase is a proton pump responsible for acidification of intracellular systems, a process necessary for the required pH-dependent fusion event between LCMV viral and cellular membranes in the acidic environment of the late-stage endosome(15, 39). To validate this screening result, known v-ATPase inhibitors Bafilomycin A_1_ (**Figure S2B**), Bafilomycin B_1_ (**Figure S2C**), and Concanamycin A (**Figure S2D**) were tested for efficacy in LCMV infection inhibition(40). As expected, all three drugs exhibited dose-dependent protection against LCMV infection in A549 cells with nanomolar efficacy, consistent with the critical role v-ATPase plays in LCMV infection.

To explore other candidate genes of interest identified in this screen, monoclonal A549 knockout (KO) cell lines containing frameshift mutations were generated for several top-scoring genes (**Figure S2E**). These cells lines were also tested for normal cell growth (**Figure S2F**). Among these candidates were the transmembrane proteins encoded by *ACKR4, CD164, EMC1, IL2RA;* the trans-Golgi/endosome membrane trafficking complex *ARFRP1* and *SYS1*; the vesicular transport associated genes *YKT6* and *RAB10*; the ZAP anti-viral protein co-factor *KHNYN*; and the signal recognition particle gene *SRP14*. In all cases, homozygous and heterozygous knockouts in A549 cells yielded significant decreases in LCMV infection, ranging from severely impaired relative to wildtype: 1.3% infected (-/-*CD164*), to moderately impaired: 77% infected (-/+*SYS1*). (**Figure 1C**)

Since knockout of CD164 demonstrated near ablation of infection, we chose to follow-up on this protein to explore its role in the viral life cycle. CD164 is a heavily glycosylated transmembrane sialomucin cell adhesion protein expressed in a wide range of tissues(41, 42). This gene was originally characterized as a marker for CD34+ hematopoietic progenitor cells where it may be involved in a variety of processes, including cellular adhesion, autophagy, tumorigenesis, and metastasis(43, 44). To date, *CD164* has not been associated with any known viral entry mechanisms.

To further investigate the role of this gene in LCMV infection, monoclonal *CD164* KO (Δ*CD164*) cell lines were generated in two additional cell types: 293T (human embryonic kidney cells) and 3T6-Swiss albino (mouse embryonic fibroblast cells). In both human lines, A549 and 293T, deletion of CD164 reduced infection by 99% and 95% respectively, while the effect in the mouse cell line 3T6 was moderate (41% reduction) (**Figure 1D**). Infectivity of each KO cell line was nearly fully restored by complementation with ectopically expressed human *CD164* (*hCD164*) gene driven by the CMV promoter. Complementation with the mouse *Cd164* (*mCD164*) gene, which is 62.32% identical on a protein level, partially restored infectivity in all three cell lines. We confirmed protein expression levels in knockout and complemented cell lines by Western blot (**Figure S2G-I**). Together, our data suggests that *CD164* is essential for LCMV infection in human cells.

### Pseudotyped viral infection shows that CD164 is a LCMV-specific mammarenavirus human entry factor

Previous work has demonstrated *DAG1* to be an entry-related attachment receptor in mice(17). Our screen also identified DAG1 as an important LCMV entry factor in addition to implicating CD164 as a determinant of human cell entry for LCMV. To further explore the dependency of mammarenaviruses on *CD164* or *DAG1* for viral entry, we generated and validated additional monoclonal KO cell lines, Δ*DAG1* and Δ*CD164*/Δ*DAG1* double KO, in both A549 (**Figure S3A**) and 293T (**Figure S3B**) cell backgrounds. To specifically test the entry stage of the viral life cycle, recombinant green fluorescent protein (GFP) expressing vesicular stomatitis virus (rVSV-ΔG(GFP)) pseudotyped with a panel of mammarenavirus GP were utilized(45). The advantage of this method is two-fold: targeted examination of GP receptor tropism in the absence of other factors that may influence native viral infection and the ability to study BSL-4 pathogenic mammarenaviruses in standard BSL-2 laboratory conditions(46).

The GPs from several LCMV strains representing a range of DAG1 affinities were combined with rVSV-ΔG(GFP) to generate pseudotyped virus **(Figure2A-E)**. Arm 53b (used for the CRISPR screen) (**Figure 2A**) and WE2.2 (**Figure 2B**) represent low DAG1 affinity strains, while Armstrong Clone 13 (Arm Cl13) (**Figure 2C**), WE54 (**Figure 2D**), and WE (**Figure 2E**) were chosen to represent high-DAG1 affinity strains(22, 23, 25). Deletion of *CD164* reduced infection by all four pseudotyped viruses by 78%-99% in both human cell lines, indicating a strong CD164 dependency in all cases. In contrast, knockout of *DAG1* in both A549 and 293T cells led to only moderate decreases in pseudotyped virus infection (23%-38% reduction for A549; up to 63% reduction for 293T) across all LCMV strains. In one case (293T Δ*DAG1* infected with rVSV-ΔG(GFP)+WE54-GP), deletion of *DAG1* had virtually no measurable impact on pseudotyped virus infection. Consistent with that, Δ*DAG1/ΔCD164* double KO cells yielded decreased pseudovirus infection similar or below those observed in *CD164* KO cells. Together, these results suggest that *CD164* is the major determinant for LCMV entry in both A549 and 293T cells, whereas *DAG1* plays only an accessory role in these human cell types.

**Figure 2.**
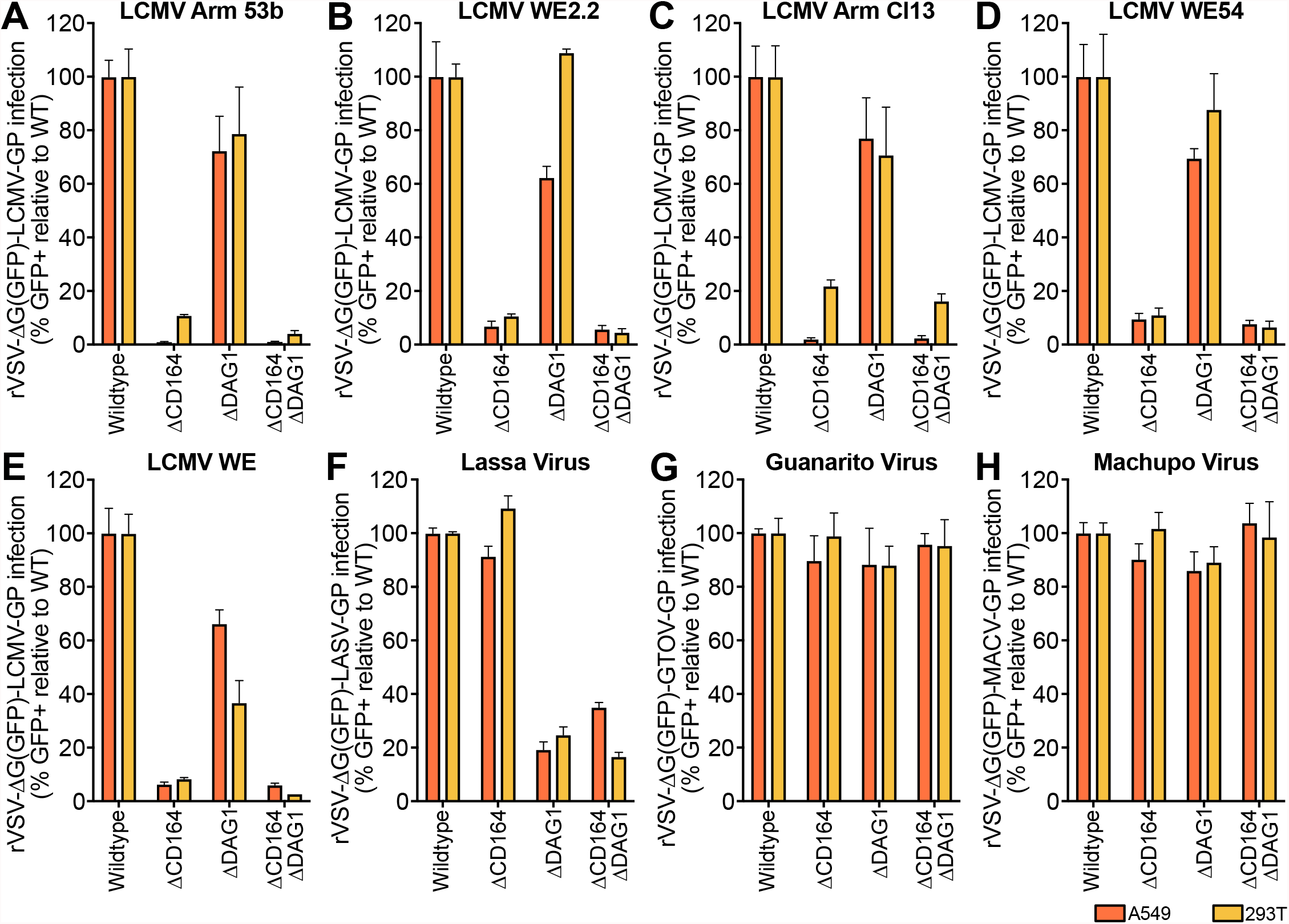
Infection of KO cell lines with a panel of mammarenavirus GP pseudotyped virus. (A-E) Percent infection of Δ*CD164*, Δ*DAG1*, and Δ*CD164*/Δ*DAG1* double-KO cells relative to WT in either A549 or 293T cell type backgrounds following inoculation with low DAG1 affinity LCMV strains (A) Armstrong 53b-GP or (B) WE2.2-GP, and high DAG1 affinity strains (C) Armstrong Clone 13-GP, (D) W54-GP, or (E) WE-GP pseudotyped virus as determined by flow cytometry for GFP positivity. Cells were infected at MOI 1 and measured 24 hpi. Error bars indicate standard error of three independent experiments. (F-H) Percent infection of ΔCD164, ΔDAG1, and ΔCD164/ΔDAG1 double-KO cells relative to WT in either A549 or 293T cell type backgrounds following inoculation with (D) LASV-GP, (E) GTOV-GP, or (F) MACV-GP pseudotyped virus as determined by flow cytometry for GFP positivity. Cells were infected at MOI 1 and measured 24 hpi. Error bars indicate standard error of three independent experiments.

To extend these findings beyond LCMV, a selection of hemorrhagic mammarenaviruses GPCs were used to generate pseudovirus for infection in A549 and 293T cells. As previously described, *DAG1* is an important entry factor for the OW mammarenavirus Lassa Virus (LASV), but not for the NW mammarenaviruses, Guanarito virus (GTOV) and Machupo virus (MACV)(17, 18). Consistent with these findings, deletion of *DAG1* abrogated LASV entry (**Figure 2F**) but had minimal effects on GTOV (**Figure 2G**) and MACV entry (**Figure 2H**). By contrast, deletion of *CD164* had no effect on infection for any of the three tested pathogenic mammarenaviruses, suggesting that *CD164* is a critical human entry factor for LCMV, but not other mammarenaviruses. Using GP-pseudotyped virus, we have determined that *CD164* plays a major functional role for LCMV entry in human cells and no other tested hemorrhagic mammarenaviruses, while *DAG1* is an important entry factor for LASV and, to a lesser extent, certain strains of LCMV.

### N-linked glycosylation within the cysteine-rich domain is critical for LCMV infection

*CD164* is a 197 amino acid type 1 integral transmembrane protein featuring a 14 amino acid intracellular tail and a 139 amino acid extracellular region that is expressed as a homodimer nearly ubiquitously throughout human tissues(47). The extracellular portion of CD164 is comprised of two mucin domains flanking a cysteine-rich domain. The protein also features one predicted attachment site for O-linked glycans and 9 predicted N-linked glycosylation sites throughout the mucin and cysteine-rich domains.

To further dissect the role of *CD164* with respect to LCMV entry, a series of *CD164* domain deletion mutants were constructed and introduced into A549 Δ*CD164* and 293T Δ*CD164* cells (**Figure 3A**). Deletion of the first mucin domain (Δ*CD164* + *hCD164*(ΔE1) did not affect infection, suggesting this domain is not necessary for LCMV entry. Extending the deletion into the cysteine-rich domain (Δ*CD164* + *hCD164*(ΔE1-2)) however, ablated infection (mean 98% reduction for A549; 86% reduction for 293T), thereby phenocopying the ΔCD164 cells. We confirmed expression of all domain deletion constructs by Western blot (**Figure S4A and S4B**).

**Figure 3.**
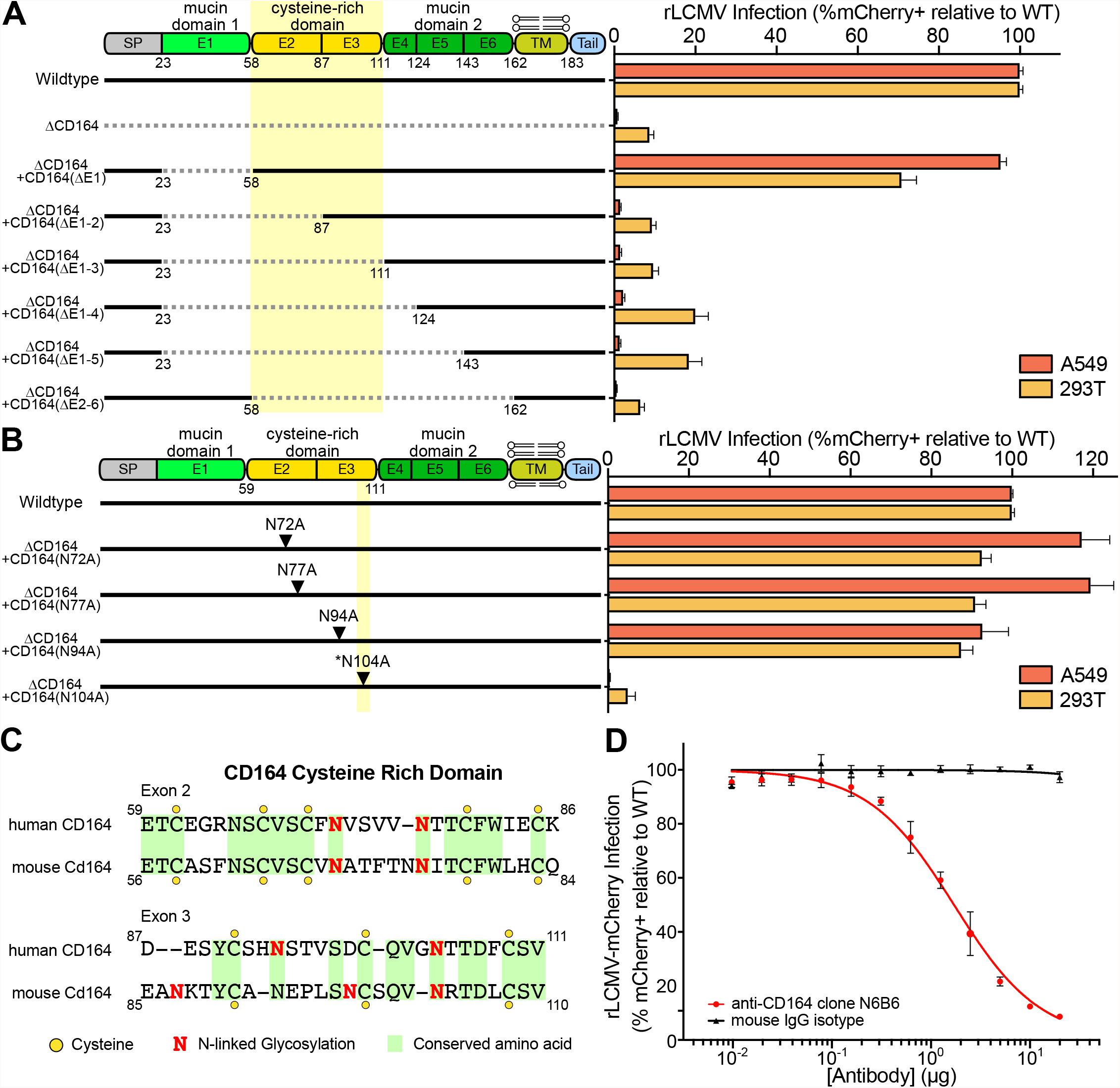
CD164 functional region determination through anti-body binding, domain deletion, and alanine mutagenesis. (A) Schematic (left) of wildtype, Δ*CD164*, Δ*CD164* + *hCD164*(ΔE1), Δ*CD164* + *hCD164*(ΔE1-2), Δ*CD164* + *hCD164*(ΔE1-3), Δ*CD164* + *hCD164*(ΔE1-4), Δ*CD164* + *hCD164*(ΔE1-5), and Δ*CD164* + *hCD164*(ΔE2-6). Complemented A549 and 293T cells were challenged with rLCMV-mCherry (MOI 1) and infection was measured by flow cytometry at 24 hpi (right). Percent infection was normalized to wildtype. Error bars represent standard error of three independent experiments. (B) Schematic (left) of wildtype, ΔCD164 KO + hCD164(N72A), ΔCD164 + hCD164(N77A), ΔCD164 + hCD164(N94A), and ΔCD164 + hCD164(N104A). Complemented A549 and 293T cells were challenged with rLCMV-mCherry (MOI 1) and infection was measured by flow cytometry at 24 hpi (right). Percent infection was normalized to wildtype. Error bars represent standard error of three independent experiments. (C) Amino acid similarities of the cysteine-rich region in human CD164 and mouse Cd164 determined using the ClustalW program on SnapGene. Yellow circles indicate cysteine residues, red N symbolizes N-linked glycosylation sites, and identical amino acids are highlighted in green. (D) Blockade of LCMV infection with serial dilutions of anti-human CD164 monoclonal mouse antibody clone N6B6 or mouse IgG2a-κ isotope control in wild type A549 cells. Cells were infected at MOI 1 and infection measured at 24 hpi. Error bars indicate standard error of three independent experiments.

The cysteine-rich region of *CD164* contains four putative N-linked glycosylation sites (**Figure 3B**). To test the importance of these sites individually, alanine substitutions were introduced in place of each relevant asparagine, and expression of these mutant construct was confirmed by Western blot (**Figure S4C and S4D**). Mutation of N-linked glycosylation sites at positions 72, 77, and 94 did not reduce infection by rLCMV-mCherry, however, substitution of N104 completely abolished infection. This asparagine residue, which is conserved between human and mouse CD164 (**Figure 3C**), sits in a loop region between a beta-sheet and an alpha-helix as predicted by AlphaFold (**Figure S4E**)(48). The ablation of infection due to mutagenesis of the N-linked glycosylation site suggests that the cysteine-rich domain, including a critical asparagine amino acid, is required for *CD164*-mediated infection by LCMV.

The deletion mapping of *CD164* indicated the importance of the cysteine-rich domain. To further explore this domain, we tested whether an anti-CD164 monoclonal antibody (mAb) could competitively inhibit LCMV infection. The anti-CD164 mAb N6B6, which was demonstrated to bind a conformationally dependent backbone epitope encompassing the cysteine-rich domain between the two mucin domains(42, 49), blocked infection by rLCMV-mCherry in a dose-dependent manner (**Figure 3D**). These results are consistent with the deletion mapping and alanine mutagenesis data, highlighting the importance of the central cysteine-rich domain for LCMV infection.

### CD164 is highly expressed in human placenta and mediates LCMV infection in placental cells

Although LCMV infection as a child or an adult are typically inconsequential, infection during pregnancy can lead to transplacental human fetal infections with severe clinical consequences(10). Like many other congenital pathogens, LCMV has tropism for fetal neural and retinal tissue, leading to developmental issues such as microencephaly, macrocephaly, chorioretinitis, periventrictular calcification, and hydrocephalus(50, 51). Retrospective studies on serologically confirmed cases show that children with congenital LCMV infection have a 35% mortality rate by 2 years of age and survivors experience long-term neurological, motor, and visual impairments(9, 52).

Human fetal vulnerability to LCMV led us to hypothesize that CD164 may play a role in transplacental infection. To explore tissue specific expression of CD164 during pregnancy, healthy second trimester placentas were co-stained with CD164 mAb anti-CD164 N6B6. CD164 was highly expressed in the outer layer of floating chorionic villi and absent in the underlying mesenchyme; co-localization with cytokeratin-7 confirmed that CD164 was expressed in cytotrophoblasts (**Figure 4A**)(53). In contrast, CD164 was not detected in the decidua (maternal side), pointing to a fetal-specific localization at this interface (**Figure S5A**). Cytotrophoblasts bathe in maternal blood and are an initial contact site for pathogens(54), suggesting that CD164 is present in tissue structures and locations amenable for transplacental infection of the developing fetus.

**Figure 4.**
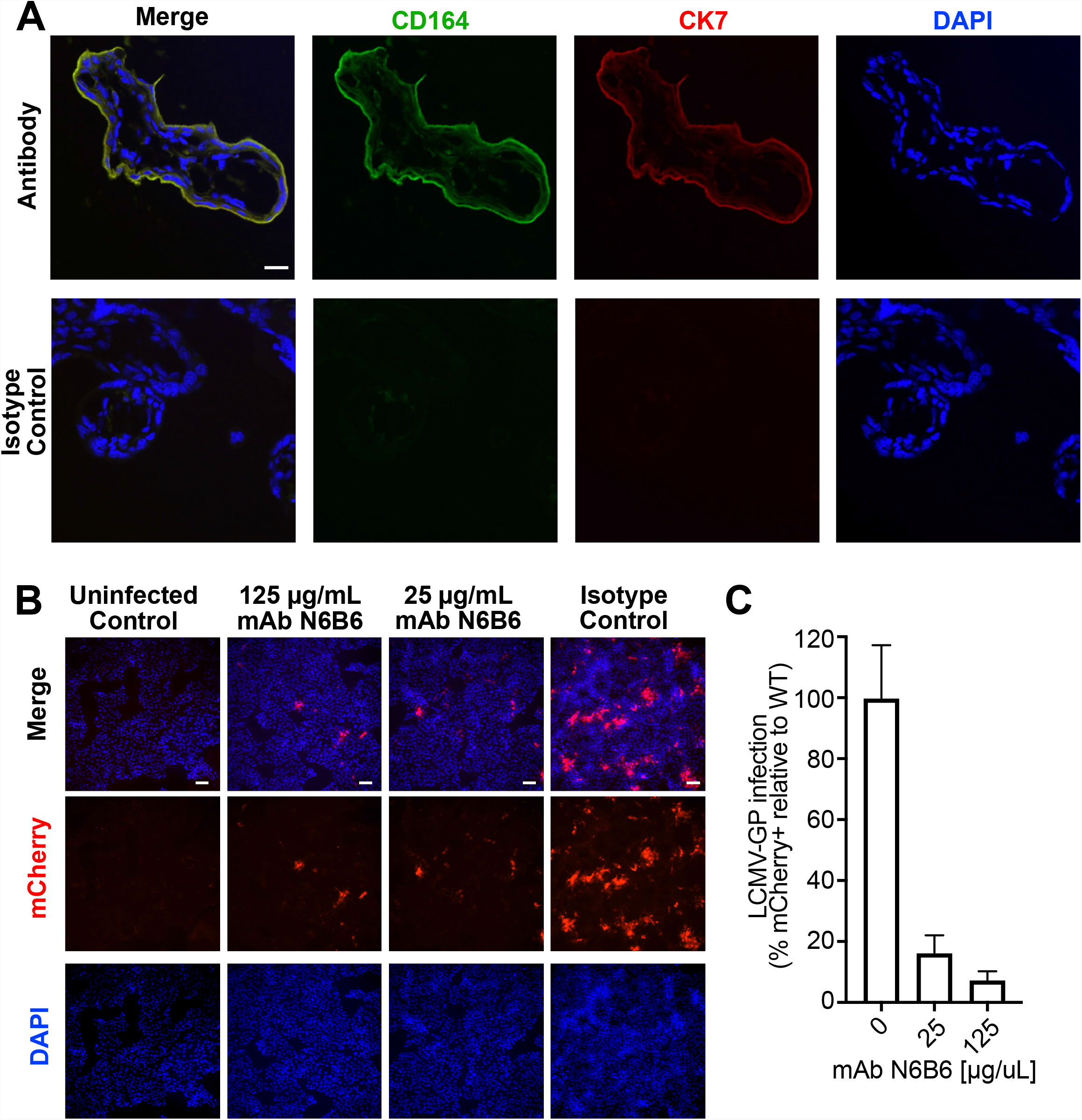
Characterization of CD164 as a therapeutic target in human placenta. (A) Double immunofluorescence staining for CD164 and CK7 or isotypes staining followed by counterstaining with DAPI in villous trophoblastic tissue. Original images were taken by confocal microscopy at 100x magnification. Scale bar represents 20 μm. (B) Immunofluorescence imaging of JEG-3 placenta cells pre-incubated with various concentrations of anti-CD164 mAb N6B6 and infected with r3LCMV-mCherry at MOI 0.5. Cells were fixed and imaged at 10x magnification 24 hpi. Scale bar represents 20 μm. (C) Quantification of percent infection of JEG-3 placenta cells pre-incubated with various concentrations of anti-CD164 mAb N6B6 and infected with r3LCMV-mCherry at MOI 0.5. Analysis was done on 4 FOV in 2 independent infections and normalized to infection control.

CD164 expression was also observed in JEG-3 human chorionic cell line, which we have demonstrated to be permissive to LCMV infection (**Figure S5B and S5C**). Preincubation of these cells with anti-CD164 mAb N6B6 blocked rLCMV-mCherry infection, with a treatment of 25ug/mL N6B6 reducing the detection of mCherry-positive cells by 84% and 125ug/mL reducing infection further by 94% (**Figure 4B and 4C**). This dose-dependent inhibition of LCMV infection indicates that LCMV utilizes CD164 ectodomains in placental cells. Thus, blocking CD164-mediated entry with a targeted antibody may be a viable therapeutic intervention for congenital LCMV.

## Discussion

In this study, we performed a genome-wide CRISPR KO screen to identify host factors important for LCMV. Our results highlight a subset of genes that appear to be shared generally among mammarenaviruses. These genes span pathways and functions such as sialic acid metabolism, heparan sulfate biosynthesis, glycosylation and Golgi trafficking, and late-stage endosome acidification(25, 28, 32). Most notably, we identified 10/24 of the v-ATPase subunits (5 in each of the V_1_ and V_0_ domains) as well as four signal recognition particle subunits. The previously described LCMV entry receptor, *DAG1*, was moderately enriched, consistent with the use of the low DAG1 affinity Arm 53b LCMV strain in this screen(22, 23).

This screen also revealed genes, most notably CD164, that have not previously linked to LCMV infection, perhaps facilitated by using a human epithelial lung cell line. We found that in the absence of CD164 in human cells LCMV infection is nearly ablated. In contrast, mouse Δ*CD164* cells yielded only moderate reduction in LCMV infection. Consistent with this, when complemented with ectopically expressed human or mouse CD164, human CD164 restored infectivity, while the mouse homologue of CD164, which is 62% identical on the protein level, only partially restored infectivity. CD164 localizes to the cell surface and late-stage endosomes, consistent with the LCMV entry route for successful infection(47). Like DAG1, it is a ubiquitously expressed cell adhesion molecule present in nearly all tested human tissue. Unlike DAG1, to which LCMV strains show a range of affinity, all five LCMV strains tested here required CD164 for infection in human cells. Deletion of DAG1 partially reduced infectivity by LCMV with some variability depending on strain. These data strongly support that human lung cells require CD164, and not necessarily DAG1, for viral infection by LCMV, whilst mouse cells appear to rely on CD164 only partially.

Further characterization of *CD164* by deletion mapping and alanine mutagenesis suggests that the cysteine-rich domain, particularly a single critical N-linked glycosylation site, is required for CD164-mediated infection. The importance of the cysteine-rich domain was reinforced by blocking using the anti-CD164 mAb N6B6, whose presence can inhibit LCMV infection in a dose-dependent manner. These data together suggest that binding by N6B6 to CD164 renders the critical interaction region inaccessible, and thus preventing LCMV infection.

While LCMV infection is generally mild among adults and children, clinical outcomes following congenital infections tend to be severe. LCMV transplacental infections are typically fatal and survivors experience long-term neurological, motor, and visual impairments(9, 10, 52). While off-brand use of ribavirin occurs in cases of LCMV infection following solid organ transplants, no current treatment procedure exists for congenital LCMV(8). We demonstrated a fetal-specific localization of CD164 to cytotrophoblasts, the placental interface bathed in maternal blood and the initial contact site with pathogens. Additionally, the mAb N6B6 also inhibits LCMV infection in placental cell lines, suggesting that this interaction could be a target for possible therapeutic intervention.

Finally, there is evidence that DAG1 is not the sole receptor used by LCMV for viral entry, and that entry can occur through many different routes depending on whether the preferred receptor DAG1 is present(17, 23). In this study, we have identified a compendium of genes important for LCMV infection in human cells, including the sialomucin CD164. We demonstrated that CD164 is an essential determinant for LCMV entry into human cells, which fills a critical gap in our understanding of human tissue infection by this virus. Whether the reliance on CD164 is unique to LCMV, or whether this entry factor is utilized by additional viruses remains unknown. Given the apparent unique dependency of LCMV on CD164, and the practical implications in its involvement in transplacental infection, further exploration of the mechanistic details by which LCMV co-opts CD164 is warranted.

## Materials and Methods

### Cell lines

A549 (ATCC), 293T (ATCC), 3T6 (ATCC), BHK-21 (ATCC), and Vero cells (ATCC) were cultured in DMEM (Gibco) supplemented with 10% fetal bovine serum (FBS, Gibco), penicillin-streptomycin-glutamine (Gibco), and HEPES (Gibco) at 37C and 5% CO_2_. JEG-3 (ATCC) were cultured in EMEM (ATCC) supplemented with 10% FBS (Gibco), penicillin-streptomycin-glutamine (Gibco), non-essential amino acids (Gibco), and sodium pyruvate (Gibco) at 37C and 5% CO_2_. All cell lines tested negative for mycoplasma contamination (Lonza).

### Virus stocks

Recombinant LCMV containing an mCherry reporter (rLCMV-mCherry) was rescued from BHK-21 cells transfected with plasmids encoding viral proteins and reporter containing recombinant genome as previously described(30). Both rLCMV-mCherry and LCMV strain ARM-4 (Gift of Michael J. Buchmeier) were propagated on BHK-21 cells. Clarified supernatant were collected 48 hpi and stored at −80C. Viral titers were determined by focus assay on Vero cells. Briefly, serial 10-fold dilutions of virus stocks were used to infect cells in 96-well plates and incubated for 24 h. Infected cells were fixed with 4% paraformaldehyde (PFA), permeabilized with 0.2% Tween-20, stained with anti-LCMV-NP antibody (1.1.3)(55) and anti-mouse secondary (Alexa Fluor 488, Thermo Fisher Scientific) followed by foci counting. Antibody details can be found in **Table S3**. All experiments with LCMV or recombinant LCMV were performed in a biosafety level 2 laboratory.

### Genome-wide CRISPR screen

A549 cells were stably lentivirally transduced with Cas9-Blast (Addgene #52962, gift from Feng Zhang) and subsequently selected using blasticidin. Next, a total of 300 million A549-Cas9 cells were then transduced with the lentiviral human GeCKO v2 library (Addgene #1000000049, gift from Feng Zhang)(29) at a multiplicity of infection (MOI) of 0.5 and selected using puromycin for 6 days. To conduct the host factor screen, 120 million (60 million each of sub-library A and B) A549-Cas9-Blast GeCKO library cells were infected with rLCMV at MOI of 10. At 24 hpi, cells that remained mCherry-negative were collected using a Sony SH800 cell sorter.

Simultaneously, 120 million cells of uninfected A549-Cas9-Blast GeCKO library cells were collected to assess sgRNA representation as a reference.

Genomic DNA (gDNA) was extracted using the NucleoSpin Blood kit (Macherey-Nagel). The sgRNA expression cassettes were amplified from gDNA in a two-step nested PCR using Q5 High-Fidelity 2X Master Mix (NEB). For PCR-I, 48 reactions (for control samples) and 12-24 reactions (for mCherry-negative sorted FACS samples) containing 1 μg were amplified for 16 cycles. Reactions were pooled, mixed and the appropriately sized amplicons were cleaned and selected for using SPRIselect (Beckman Coulter). During PCR-II, 10 reactions containing 5 μL of PCR-I product were amplified for 10 cycles using indexed Illumina primers. PCR products were cleaned using AmpureXP beads (Beckman Coulter) and sequenced on an Illumina NextSeq 500 using a custom sequencing primer. Primer sequences can be found in **Table S4**.

Demultiplexed FASTQ files were aligned to a reference table containing sgRNA sequences and the abundance of each sgRNA was determined for each starting and sorted cell population. Guide count tables were further processed using MAGeCK to determine positive enrichment scores for each gene(31). Gene ontology enrichment was determined with statistical overrepresentation test on PANTHER(38) using genes from the 300 highest MAGeCK scores.

### Generation of monoclonal KO cell lines

sgRNA sequences against gene targets were designed using CRISPick(56) and the corresponding DNA oligos (IDT) were annealed and ligated into pX458 (Addgene #48138, gift from Feng Zhang)(57). Cells were transfected with pX458 constructs using *Trans*IT-X2 (Mirus Bio) and GFP positive cells were sorted into 96-well plates using a FACSAria II (BD) two days later. Clonal populations were genotyped by Sanger sequencing the PCR amplified sgRNA-targeted sites in the gDNA extracted using DNA QuickExtract (Lucigen). Resulting sequences were compared to references and clones containing a frameshift indel or *de novo* stop codon were selected. To determine cell growth of A549 WT and KO cell lines, CellTiter-Glo (Promega) was mixed 1:1 with cells seeded in 96-well plates for three consecutive days and the luminescence signal was quantified using the GloMax-Multi microplate reader (Promega). A list of all used sgRNA sequences and genotyping primers can be found in **Table S4**.

### Plasmids, cloning, and lentivirus production

Human CD164 (Origene, #RC202234) and mouse Cd164 (Origene, #MR201951) cDNAs were cloned into EcoRV-cut plenti-CMV-Puro-DEST (Addgene #17452, gift from Eric Campeau & Paul Kaufman)(58) using NEBuilder HiFi DNA Assembly Master Mix (NEB). Primers used to assemble expression plasmids for domain deletion mapping and alanine scanning mutagenesis of CD164 can be found in **Table S4**.

Lentivirus was produced in HEK293T by co-transfection of cDNA containing lentiviral plasmid together with helper plasmids pMD2.G (Addgene #12259, gift from Didier Trono) and pCMV-dR8.91 (Life Science Market) using *Trans*IT-Lenti (Mirus Bio). Supernatant were collected 48 h post-transfection, filtered, and added to recipient cells in the presence of Polybrene (EMD Millipore). Transduced cells were subsequently selected using Puromycin (Thermo Fisher Scientific) during days 3-5.

### Compound inhibition and antibody neutralization

Bafilomycin A_1_, Bafilomycin B_1_ (Cayman Chemical Company), and Concanamycin A (Santa Cruz Biotechnology) were resuspended in DMSO and stored at −20C until use. Cells were incubated with compounds for 1 h at 37C prior to infection assay.

Antibody neutralization assays were conducted by pre-incubating cells with anti-CD164 clone N6B6 (BD Pharmingen) or mouse IgG isotype control (BD Pharmingen) for 1 h at 37C prior to infection assay. Antibody details can be found in **Table S3**.

### Generation of Arenavirus pseudotyped vesicular stomatitis virus

Glycoprotein from LASV (Genbank: AAA46286.1), GTOV (Genbank: AAN05423.1), MACV (Genbank: AIG51558.1), and LCMV strain WE-HPI (Addgene #15793, gift from Miguel Sena-Esteves)(59) were cloned into a pCAGGS vector backbone using NEBuilder HiFi DNA Assembly Master Mix (NEB). To generate an LCMV strain Cl13-GP (Genbank: DQ361065.2) expression plasmid, mutations N176D and F260L were introduced to pCAGGS-LCMV-Arm4-GP using site-directed mutagenesis. To generate LCMV strain WE54-GP (Genbank: AJ297484.1), mutations V94A, S133T, Y155H, and T211A were introduced into LCMV strain WE-HPI-GP. To generate LCMV strain WE2.2-GP (Genbank: AJ318512.1), mutation S153F was introduced into LCMV strain WE54-GP(22, 23). A list of primers used for cloning and site-directed mutagenesis can be found in **Table S4**.

To rescue the various VSV-ΔG-Arenavirus-GP pseudotype virus, 293T cells were transfected with arenavirus glycoprotein expression plasmids using *Trans*IT-LT1 (Mirus Bio). Cells were transduced the following day with VSV-ΔG-GFP (Kerafast)(45) at MOI 3 and incubated in media containing anti-VSV-G antibodies (Kerafest) for 24 h. Clarified supernatant containing pseudovirus were collected and stored at −80C. Stock titers were measured using flow cytometry on a FACSCelesta (BD). All experiments with pseudotyped VSV were performed in a biosafety level 2 laboratory.

### Flow cytometry analysis of viral infection assays

Cells plated in 96-well plates were infected with rLCMV-mCherry or LCMV at MOI 1 for an adsorption period of 1 h at 37C and subsequently cultured for 24 h. To analyze percent infected, cells were trypsinized and fixed in suspension with 4% PFA for 30 min. For infection with rLCMV-mCherry, analysis was done by flow cytometry on FACSCelesta (BD) where approximately 5,000 cells were recorded and gated based on SFC/SSC, FSC-H/FSC-A (singlets), and PE-CF594 (mCherry) using FlowJo 10. For infection with LCMV, cells were permeabilized and stained for LCMV-N protein (primary: 1.1.3, secondary: Alexa Fluor 488) prior to flow analysis, with gating for FITC (eGFP). Antibody details can be found in **Table S3**.

For pseudotype infection assays, cells seeded in a 96-well plate were infected with various VSV-Arenavirus-GP pseudoviruses. At 24 hpi, cells were lifted using Tryple Select Enzyme (Gibco) and flowed on a FACSCelesta (BD), and for FITC (eGFP) signal as previously described.

### Western Blots

Cells were scraped and lysed in RIPA buffer on ice for 30 min. All lysates were separated by SDS-PAGE on pre-cast 4-12% Bis-Tris gels (Thermo Fisher Scientific) in the NuPAGE electrophoresis system. Proteins were transferred onto nitrocellulose membrane using the Bio-Rad Mini-Protean Mini Trans-Blot transfer system. Membranes were blocked with Tris-buffered saline with 0.05% Tween-20 and 5% non-fat milk. Blocked membranes were incubated with primary antibody diluted in blocking buffer overnight at 4C on a shaker. Primary antibodies were detected by incubating membranes with 1:15,000 dilution of IRDye secondary antibodies (LI-COR) for 1 h at room temperature and visualized using the Odyssey CLx (LI-COR). Antibody details can be found in **Table S3**.

### Quantification and Statistical Analysis

Enrichment scores, p-values, and false discovery rates for the CRISPR screen were determined using the MAGeCK algorithm(31). For GO analysis, p-values were determined using Fisher’s exact test on PANTHER’s statistical overrepresentation test(38). For viral infection, drug treatment, antibody neutralization, and cell growth experiments, biological replicates are defined as independent treatments and measurements from cells harvested from multiple wells on different days. Replicates are displayed as mean ± SEM and visualized using GraphPad Prism 9. Dose-response curves for drug treatments and antibody neutralizations were generated by applying a non-linear curve fit with least-squares regression and default parameters using GraphPad Prism 9. No additional statistical tests were performed. No methods were used to determine sample size estimation or whether the data met assumptions of the statistical approaches. For all experiments, the statistical details can be found in the figure legends.

## Acknowledgements

We would like to thank Dr. Stephanie Gaw for gifting placental specimens used in our microscopy studies. This work was supported by the National Institutes of Health [F31AI150007 to S.S.]; the Chan Zuckerberg Biohub [to J.D., C.Y.W., and A.P.]; and the Chan Zuckerberg Initiative [to J.D.]. The funders had no role in study design, data collection and interpretation, or the decision to submit the work for publication.

---

**Supplementary Figure 1.**
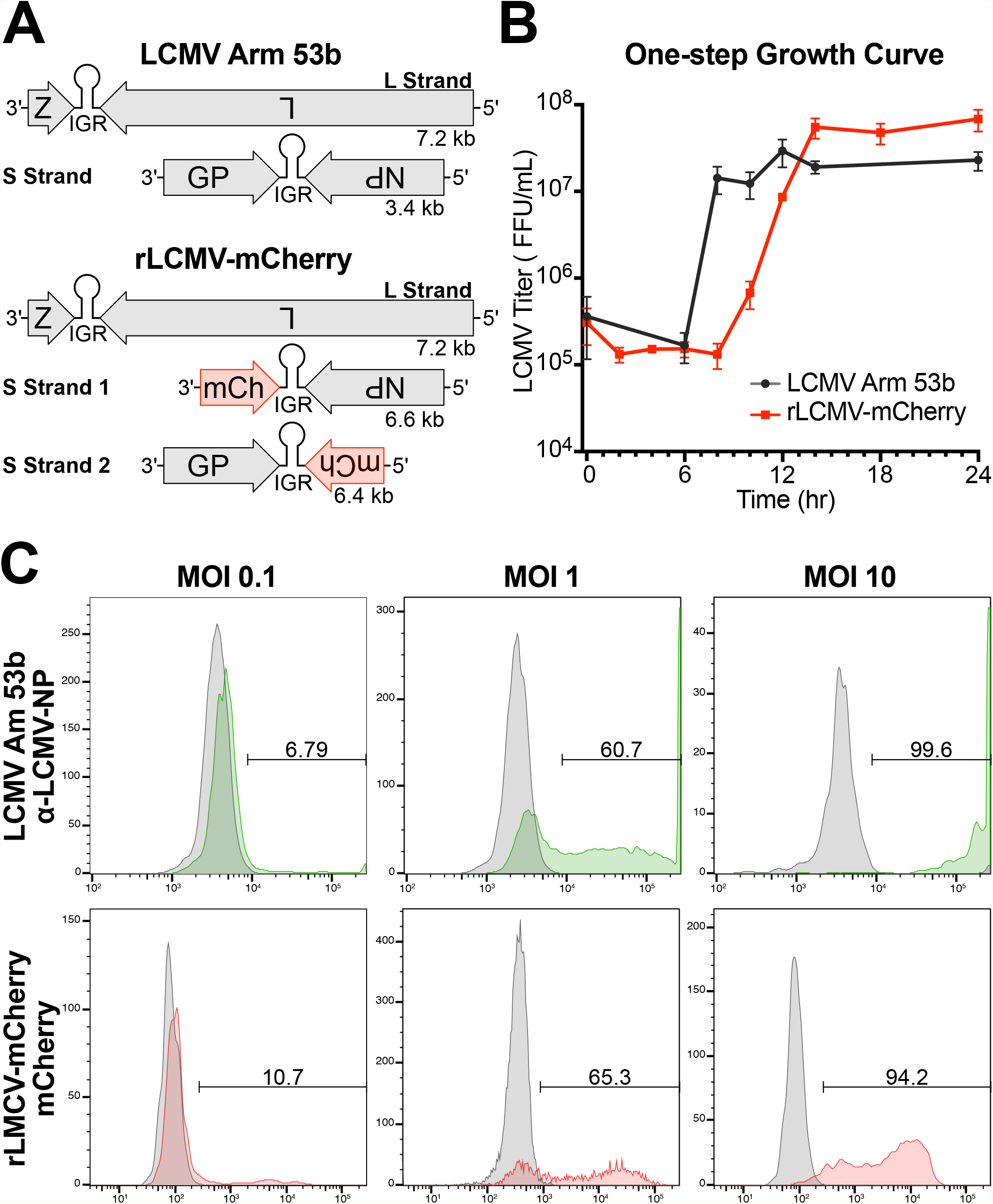
Validation of recombinant virus rLCMV-mCherry infectivity. (A) Schematic representation of LCMV Arm 53b and r3LCMV-mCherry genomes. (B) One-step growth curves of wildtype LCMV Arm 53b strain (black) and rLCMV-mCherry made in Arm 53b background (red) as measured by TCID50 over a 24-hour time course. Error bars indicate standard error of three independent experiments. (C) Infection percentage of A549 cells infected at multiplicity of infection (MOI) 0.1, 1, and 10 with wildtype LCMV Arm 53b (top) or r3LCMV-mCherry (bottom) as measured at 24 hours post infection (hpi) using flow cytometry. Cells infected with Arm 53b were stained with anti-LCMV-NP monoclonal antibody (mAb) 113 primary and Alexa 488 secondary and measured for FITC signal in comparison with an uninfected control. Cells infected with rLCMV-mCherry were measured for PE-CF594 (mCherry) signal in comparison with an uninfected control.

**Supplementary Figure 2.**
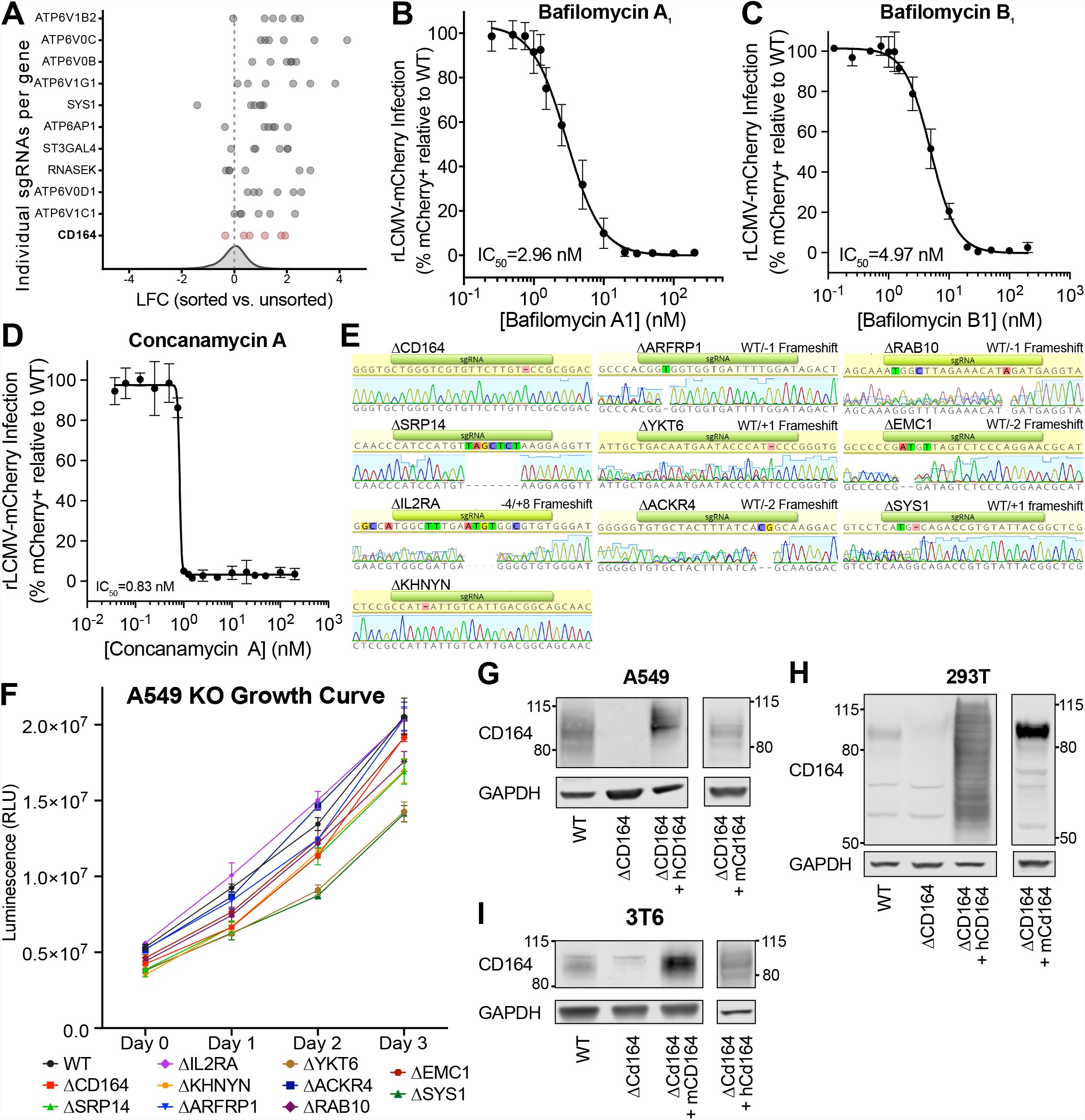
Additional hit validation and characterization of gene-edited cells. (A) Log fold changes (LFC) of individual sgRNA of the top 10 scoring genes and CD164 (red) when comparing the infected and sorted cell population versus the uninfected cell population. Overall sgRNA distribution is shown at the bottom of the graph and dotted line indicates mean LFC of all sgRNAs. (B-D) Dose-response curve of v-ATPase inhibitors on rLCMV-mCherry infection at MOI 1 in A549 cells at 24 hpi, yielding (B) Bafilomycin A_1_ IC50 = 2.96 nM, (C) Bafilomycin B_1_ IC50 = 4.97 nM, and (D) Concanamycin A IC50 = 0.83 nM. Error bars indicate standard error of three independent experiments. (E) Genotyping of clonal A549 where the target loci were PCR-amplified, Sanger-sequenced, and aligned to WT reference sequence. (F) Analysis of cell proliferation of WT and clonal A549 KO cells. Cells were plated in 96-well and proliferation was measured daily using Cell Titer Glo. Error bars indicate standard error from three separate well per cell line per time point. (G-I) Western blot analysis of WT, Δ*CD164*, Δ*CD164* + *hCD164*, and Δ*CD164* + *mCd164* for A549, 293T, and 3T6 cell lines. Human cell lines (A549 and 293T) were probed with anti-hCD164 antibody except for the *mCd164* addback which was probed with anti-mCd164 antibody. Mouse cell line 3T6 was probed with anti-mCd164 antibody except for the *hCD164* addback, which was probed with anti-hCD164 antibody. GAPDH was used as loading control.

**Supplementary Figure 3.**
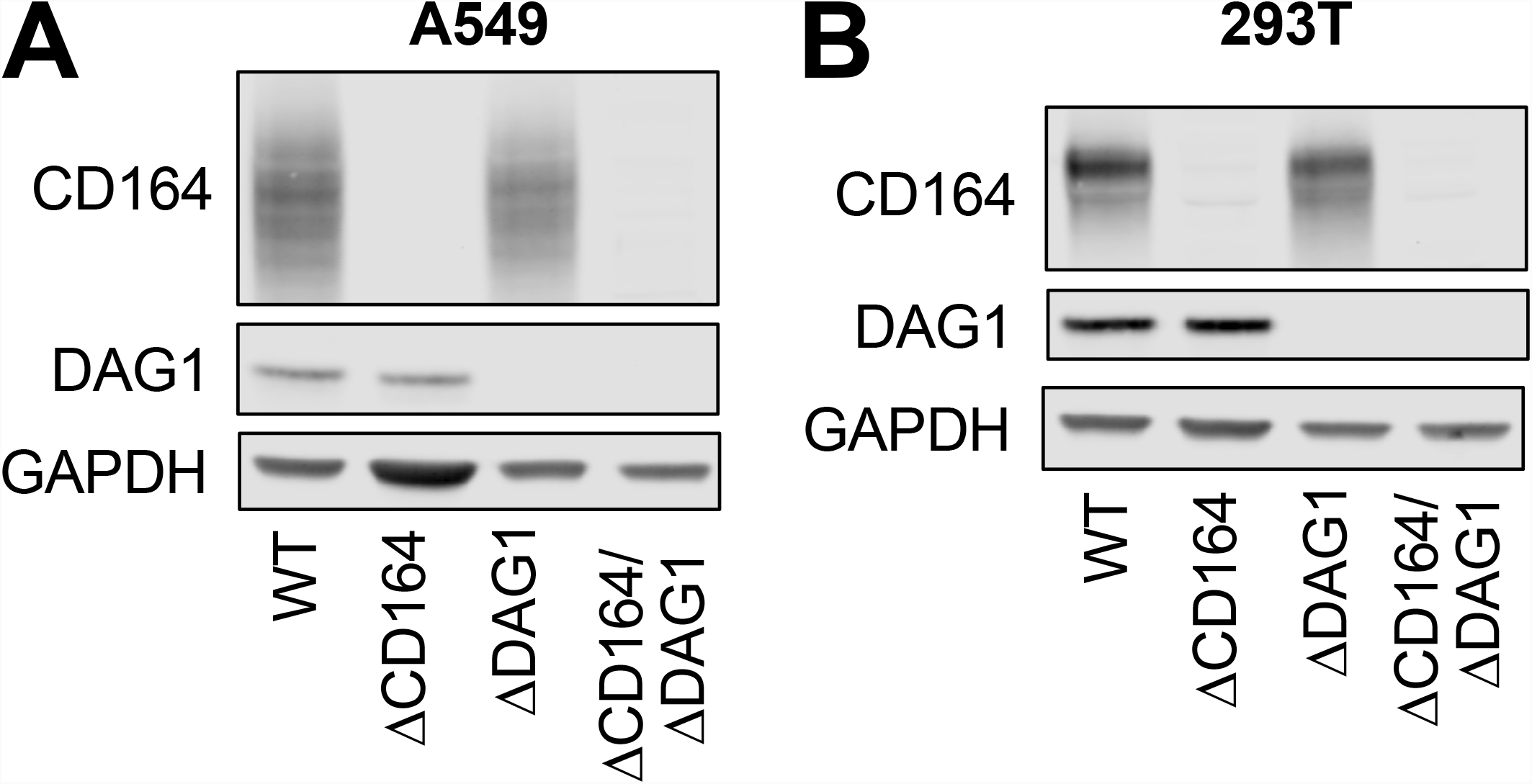
Characterization of Δ*CD164*, Δ*DAG1*, and Δ*CD164*/Δ*DAG1* double KO cells. (A-B) Western blot analysis of Δ*CD164*, Δ*DAG1*, and Δ*CD164*/Δ*DAG1* double KO cells in (A) A549 or (B) 293T cell backgrounds. GAPDH was used as a loading control.

**Supplementary Figure 4.**
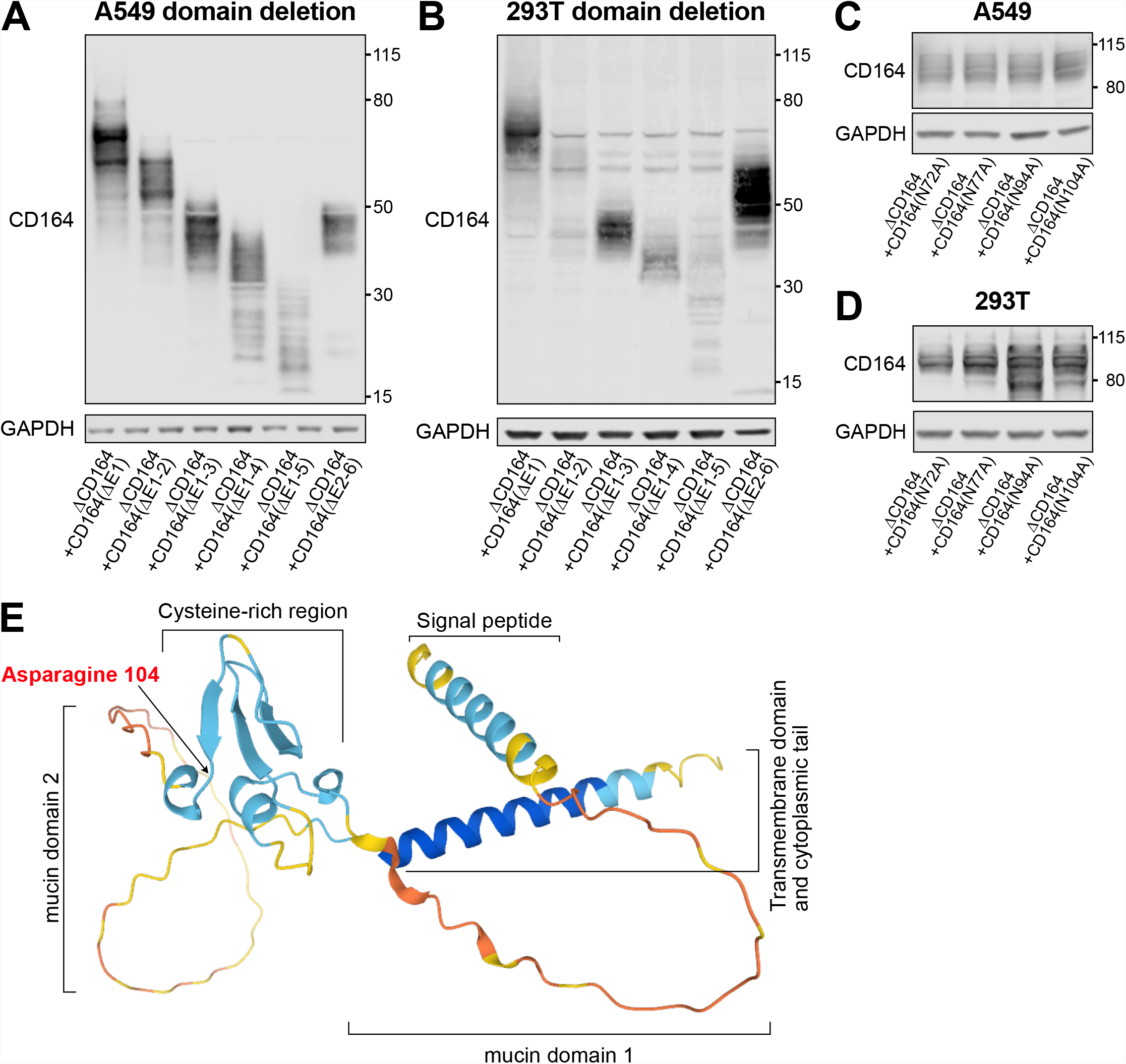
Characterization of CD164 domain deletion and alanine mutagenesis add backs. (A-B) Western blot analysis of deletion domain addbacks in (A) A549 or (B) 293T cell backgrounds. (C-D) Western blot analysis of alanine mutagenesis addbacks in (C) A549 or (D) 293T cell backgrounds. All CD164 addbacks were probed with anti-FLAG antibody. GAPDH was used as a loading control. (E) AlphaFold prediction of CD164 protein structure. Prediction had low position error for the signal peptide, the cysteine-rich region, the transmembrane domain, and the cytoplasmic tail and high position error for the two mucin domains. Location of residue 104 is noted with an arrow.

**Supplementary Figure 5.**
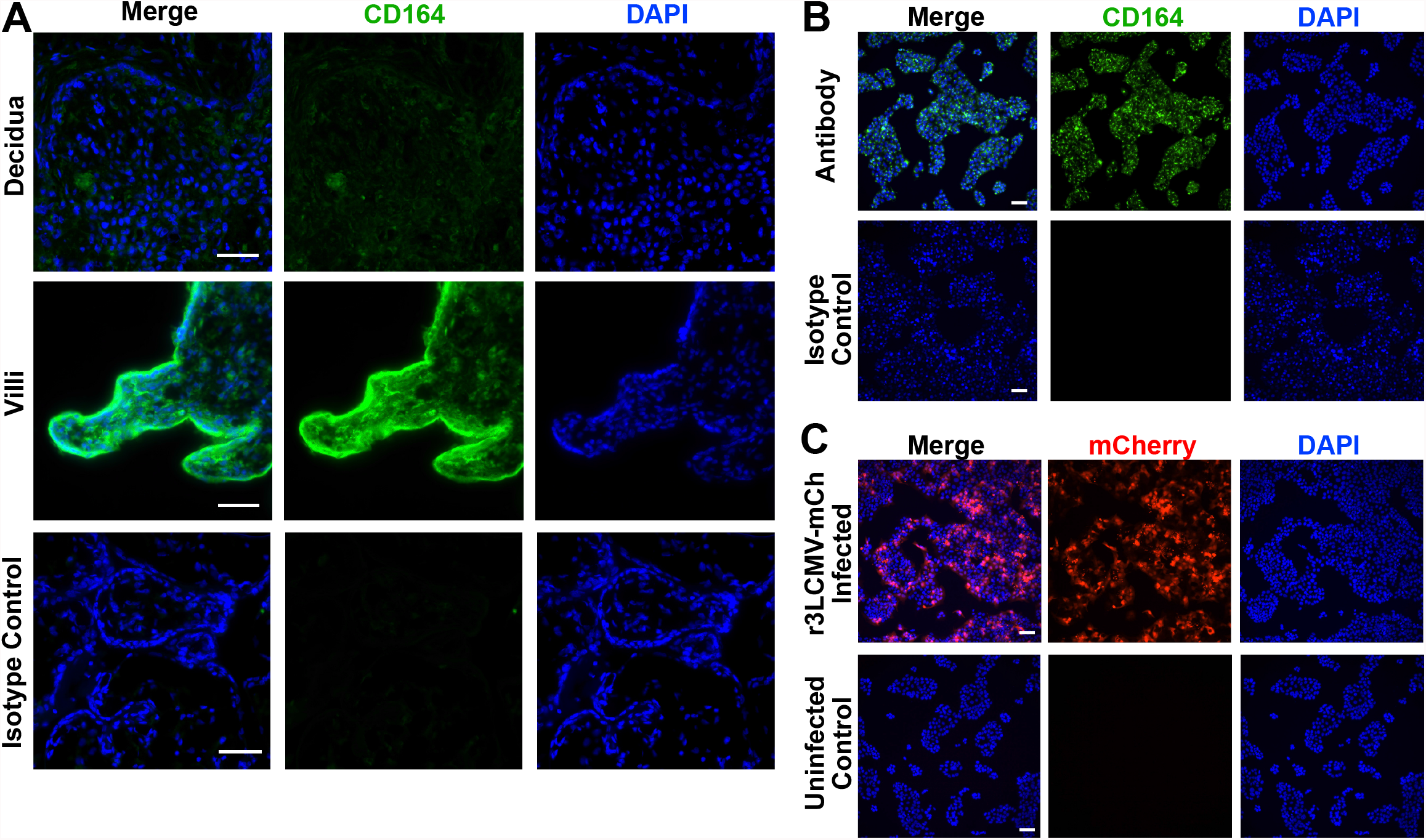
Characterization of CD164 in placenta tissue and cell line. (A) Immunofluorescence staining of CD164 or isotype control followed by counterstaining with DAPI on placenta tissue at the maternal decidua and fetal villi. Original images taken at 40x magnification. Scale bar represents 50 μm. (B) Immunofluorescence imaging of CD164 or isotype control followed by counterstaining with DAPI on JEG-3 placenta cell line. Original images taken at 10x magnification. Scale bar represents 100 μm. (C) Immunofluorescence imaging JEG-3 placenta cell line with and without infection by r3LCMV-mCherry at MOI 1 and imaged at 24 hpi. Original images taken at 10x magnification. Scale bar represents 100 μm.

## Reference

1. Radoshitzky SR, Buchmeier MJ, Charrel RN, Clegg JCS, Gonzalez J-PJ, Günther S, Hepojoki J, Kuhn JH, Lukashevich IS, Romanowski V, Salvato MS, Sironi M, Stenglein MD, de la Torre JC, Ictv Report Consortium null. 2019. ICTV Virus Taxonomy Profile: Arenaviridae. J Gen Virol 100:1200–1201.

2. Stenglein MD, Sanchez-Migallon Guzman D, Garcia VE, Layton ML, Hoon-Hanks LL, Boback SM, Keel MK, Drazenovich T, Hawkins MG, DeRisi JL. 2017. Differential Disease Susceptibilities in Experimentally Reptarenavirus-Infected Boa Constrictors and Ball Pythons. J Virol 91:e00451–17.

3. Hepojoki J, Hepojoki S, Smura T, Szirovicza L, Dervas E, Prähauser B, Nufer L, Schraner EM, Vapalahti O, Kipar A, Hetzel U. 2018. Characterization of Haartman Institute snake virus-1 (HISV-1) and HISV-like viruses-The representatives of genus Hartmanivirus, family Arenaviridae. PLoS Pathog 14:e1007415.

4. Shi M, Lin X-D, Chen X, Tian J-H, Chen L-J, Li K, Wang W, Eden J-S, Shen J-J, Liu L, Holmes EC, Zhang Y-Z. 2018. The evolutionary history of vertebrate RNA viruses. Nature 556:197–202.

5. Burri DJ, da Palma JR, Seidah NG, Zanotti G, Cendron L, Pasquato A, Kunz S. 2013. Differential recognition of Old World and New World arenavirus envelope glycoproteins by subtilisin kexin isozyme 1 (SKI-1)/site 1 protease (S1P). J Virol 87:6406–6414.

6. Traub E. 1935. A FILTERABLE VIRUS RECOVERED FROM WHITE MICE. Science 81:298–299.

7. Childs JE, Glass GE, Korch GW, Ksiazek TG, Leduc JW. 1992. Lymphocytic choriomeningitis virus infection and house mouse (Mus musculus) distribution in urban Baltimore. Am J Trop Med Hyg 47:27–34.

8. Centers for Disease Control and Prevention (CDC). 2008. Brief report: Lymphocytic choriomeningitis virus transmitted through solid organ transplantation--Massachusetts, 2008. MMWR Morb Mortal Wkly Rep 57:799–801.

9. Wright R, Johnson D, Neumann M, Ksiazek TG, Rollin P, Keech RV, Bonthius DJ, Hitchon P, Grose CF, Bell WE, Bale JF. 1997. Congenital lymphocytic choriomeningitis virus syndrome: a disease that mimics congenital toxoplasmosis or Cytomegalovirus infection. Pediatrics 100:E9.

10. Barton LL, Mets MB, Beauchamp CL. 2002. Lymphocytic choriomeningitis virus: emerging fetal teratogen. Am J Obstet Gynecol 187:1715–1716.

11. Meyer BJ, de la Torre JC, Southern PJ. 2002. Arenaviruses: genomic RNAs, transcription, and replication. Curr Top Microbiol Immunol 262:139–157.

12. Beyer WR, Pöpplau D, Garten W, von Laer D, Lenz O. 2003. Endoproteolytic processing of the lymphocytic choriomeningitis virus glycoprotein by the subtilase SKI-1/S1P. J Virol 77:2866–2872.

13. Borrow P, Oldstone MB. 1992. Characterization of lymphocytic choriomeningitis virus-binding protein(s): a candidate cellular receptor for the virus. J Virol 66:7270–7281.

14. Eschli B, Quirin K, Wepf A, Weber J, Zinkernagel R, Hengartner H. 2006. Identification of an N-terminal trimeric coiled-coil core within arenavirus glycoprotein 2 permits assignment to class I viral fusion proteins. J Virol 80:5897–5907.

15. Di Simone C, Zandonatti MA, Buchmeier MJ. 1994. Acidic pH triggers LCMV membrane fusion activity and conformational change in the glycoprotein spike. Virology 198:455–465.

16. Burns JW, Buchmeier MJ. 1991. Protein-protein interactions in lymphocytic choriomeningitis virus. Virology 183:620–629.

17. Cao W, Henry MD, Borrow P, Yamada H, Elder JH, Ravkov EV, Nichol ST, Compans RW, Campbell KP, Oldstone MB. 1998. Identification of alpha-dystroglycan as a receptor for lymphocytic choriomeningitis virus and Lassa fever virus. Science 282:2079–2081.

18. Spiropoulou CF, Kunz S, Rollin PE, Campbell KP, Oldstone MBA. 2002. New World arenavirus clade C, but not clade A and B viruses, utilizes alpha-dystroglycan as its major receptor. J Virol 76:5140–5146.

19. Holt KH, Crosbie RH, Venzke DP, Campbell KP. 2000. Biosynthesis of dystroglycan: processing of a precursor propeptide. FEBS Lett 468:79–83.

20. Kunz S, Rojek JM, Kanagawa M, Spiropoulou CF, Barresi R, Campbell KP, Oldstone MBA. 2005. Posttranslational modification of alpha-dystroglycan, the cellular receptor for arenaviruses, by the glycosyltransferase LARGE is critical for virus binding. J Virol 79:14282–14296.

21. Rojek JM, Spiropoulou CF, Campbell KP, Kunz S. 2007. Old World and clade C New World arenaviruses mimic the molecular mechanism of receptor recognition used by alpha-dystroglycan’s host-derived ligands. J Virol 81:5685–5695.

22. Smelt SC, Borrow P, Kunz S, Cao W, Tishon A, Lewicki H, Campbell KP, Oldstone MB. 2001. Differences in affinity of binding of lymphocytic choriomeningitis virus strains to the cellular receptor alpha-dystroglycan correlate with viral tropism and disease kinetics. J Virol 75:448–457.

23. Kunz S, Sevilla N, Rojek JM, Oldstone MBA. 2004. Use of alternative receptors different than alpha-dystroglycan by selected isolates of lymphocytic choriomeningitis virus. Virology 325:432–445.

24. Shimojima M, Kawaoka Y. 2012. Cell surface molecules involved in infection mediated by lymphocytic choriomeningitis virus glycoprotein. J Vet Med Sci 74:1363–1366.

25. Volland A, Lohmüller M, Heilmann E, Kimpel J, Herzog S, von Laer D. 2021. Heparan sulfate proteoglycans serve as alternative receptors for low affinity LCMV variants. PLoS Pathog 17:e1009996.

26. Brouillette RB, Phillips EK, Patel R, Mahauad-Fernandez W, Moller-Tank S, Rogers KJ, Dillard JA, Cooney AL, Martinez-Sobrido L, Okeoma C, Maury W. 2018. TIM-1 Mediates Dystroglycan-Independent Entry of Lassa Virus. J Virol 92:e00093–18.

27. Jae LT, Raaben M, Herbert AS, Kuehne AI, Wirchnianski AS, Soh TK, Stubbs SH, Janssen H, Damme M, Saftig P, Whelan SP, Dye JM, Brummelkamp TR. 2014. Virus entry. Lassa virus entry requires a trigger-induced receptor switch. Science 344:1506–1510.

28. Raaben M, Jae LT, Herbert AS, Kuehne AI, Stubbs SH, Chou Y-Y, Blomen VA, Kirchhausen T, Dye JM, Brummelkamp TR, Whelan SP. 2017. NRP2 and CD63 Are Host Factors for Lujo Virus Cell Entry. Cell Host Microbe 22:688–696.e5.

29. Sanjana NE, Shalem O, Zhang F. 2014. Improved vectors and genome-wide libraries for CRISPR screening. Nat Methods 11:783–784.

30. Emonet SF, Garidou L, McGavern DB, de la Torre JC. 2009. Generation of recombinant lymphocytic choriomeningitis viruses with trisegmented genomes stably expressing two additional genes of interest. Proc Natl Acad Sci U S A 106:3473–3478.

31. Li W, Xu H, Xiao T, Cong L, Love MI, Zhang F, Irizarry RA, Liu JS, Brown M, Liu XS. 2014. MAGeCK enables robust identification of essential genes from genome-scale CRISPR/Cas9 knockout screens. Genome Biol 15:554.

32. Jae LT, Raaben M, Riemersma M, van Beusekom E, Blomen VA, Velds A, Kerkhoven RM, Carette JE, Topaloglu H, Meinecke P, Wessels MW, Lefeber DJ, Whelan SP, van Bokhoven H, Brummelkamp TR. 2013. Deciphering the glycosylome of dystroglycanopathies using haploid screens for lassa virus entry. Science 340:479–483.

33. Shin H-W, Kobayashi H, Kitamura M, Waguri S, Suganuma T, Uchiyama Y, Nakayama K. 2005. Roles of ARFRP1 (ADP-ribosylation factor-related protein 1) in post-Golgi membrane trafficking. J Cell Sci 118:4039–4048.

34. Zhang T, Hong W. 2001. Ykt6 forms a SNARE complex with syntaxin 5, GS28, and Bet1 and participates in a late stage in endoplasmic reticulum-Golgi transport. J Biol Chem 276:27480–27487.

35. Khoury G, Lee MY, Ramarathinam SH, McMahon J, Purcell AW, Sonza S, Lewin SR, Purcell DFJ. 2021. The RNA-Binding Proteins SRP14 and HMGB3 Control HIV-1 Tat mRNA Processing and Translation During HIV-1 Latency. Front Genet 12:680725.

36. Kisaka JK, Ratner L, Kyei GB. 2020. The Dual-Specificity Kinase DYRK1A Modulates the Levels of Cyclin L2 To Control HIV Replication in Macrophages. J Virol 94:e01583–19.

37. Tsunetsugu-Yokota Y, Honda M. 1990. Effect of cytokines on HIV release and IL-2 receptor alpha expression in monocytic cell lines. J Acquir Immune Defic Syndr (1988) 3:511–516.

38. Mi H, Ebert D, Muruganujan A, Mills C, Albou L-P, Mushayamaha T, Thomas PD. 2021. PANTHER version 16: a revised family classification, tree-based classification tool, enhancer regions and extensive API. Nucleic Acids Res 49:D394–D403.

39. Forgac M. 2007. Vacuolar ATPases: rotary proton pumps in physiology and pathophysiology. Nat Rev Mol Cell Biol 8:917–929.

40. Dröse S, Bindseil KU, Bowman EJ, Siebers A, Zeeck A, Altendorf K. 1993. Inhibitory effect of modified bafilomycins and concanamycins on P-and V-type adenosinetriphosphatases. Biochemistry 32:3902–3906.

41. Uhlén M, Fagerberg L, Hallström BM, Lindskog C, Oksvold P, Mardinoglu A, Sivertsson Å, Kampf C, Sjöstedt E, Asplund A, Olsson I, Edlund K, Lundberg E, Navani S, Szigyarto CA-K, Odeberg J, Djureinovic D, Takanen JO, Hober S, Alm T, Edqvist P-H, Berling H, Tegel H, Mulder J, Rockberg J, Nilsson P, Schwenk JM, Hamsten M, von Feilitzen K, Forsberg M, Persson L, Johansson F, Zwahlen M, von Heijne G, Nielsen J, Pontén F. 2015. Proteomics. Tissue-based map of the human proteome. Science 347:1260419.

42. Watt SM, Bühring HJ, Rappold I, Chan JY, Lee-Prudhoe J, Jones T, Zannettino AC, Simmons PJ, Doyonnas R, Sheer D, Butler LH. 1998. CD164, a novel sialomucin on CD34(+) and erythroid subsets, is located on human chromosome 6q21. Blood 92:849–866.

43. Zannettino AC, Bühring HJ, Niutta S, Watt SM, Benton MA, Simmons PJ. 1998. The sialomucin CD164 (MGC-24v) is an adhesive glycoprotein expressed by human hematopoietic progenitors and bone marrow stromal cells that serves as a potent negative regulator of hematopoiesis. Blood 92:2613–2628.

44. Havens AM, Jung Y, Sun YX, Wang J, Shah RB, Bühring HJ, Pienta KJ, Taichman RS. 2006. The role of sialomucin CD164 (MGC-24v or endolyn) in prostate cancer metastasis. BMC Cancer 6:195.

45. Whitt MA. 2010. Generation of VSV pseudotypes using recombinant ΔG-VSV for studies on virus entry, identification of entry inhibitors, and immune responses to vaccines. J Virol Methods 169:365–374.

46. Fukushi S, Tani H, Yoshikawa T, Saijo M, Morikawa S. 2012. Serological assays based on recombinant viral proteins for the diagnosis of arenavirus hemorrhagic fevers. Viruses 4:2097–2114.

47. Chan JY, Lee-Prudhoe JE, Jorgensen B, Ihrke G, Doyonnas R, Zannettino AC, Buckle VJ, Ward CJ, Simmons PJ, Watt SM. 2001. Relationship between novel isoforms, functionally important domains, and subcellular distribution of CD164/endolyn. J Biol Chem 276:2139–2152.

48. Varadi M, Anyango S, Deshpande M, Nair S, Natassia C, Yordanova G, Yuan D, Stroe O, Wood G, Laydon A, Žídek A, Green T, Tunyasuvunakool K, Petersen S, Jumper J, Clancy E, Green R, Vora A, Lutfi M, Figurnov M, Cowie A, Hobbs N, Kohli P, Kleywegt G, Birney E, Hassabis D, Velankar S. 2022. AlphaFold Protein Structure Database: massively expanding the structural coverage of protein-sequence space with high-accuracy models. Nucleic Acids Res 50:D439–D444.

49. Doyonnas R, Yi-Hsin Chan J, Butler LH, Rappold I, Lee-Prudhoe JE, Zannettino AC, Simmons PJ, Bühring HJ, Levesque JP, Watt SM. 2000. CD164 monoclonal antibodies that block hemopoietic progenitor cell adhesion and proliferation interact with the first mucin domain of the CD164 receptor. J Immunol 165:840–851.

50. Jamieson DJ, Kourtis AP, Bell M, Rasmussen SA. 2006. Lymphocytic choriomeningitis virus: an emerging obstetric pathogen? Am J Obstet Gynecol 194:1532–1536.

51. Bonthius DJ. 2012. Lymphocytic choriomeningitis virus: an underrecognized cause of neurologic disease in the fetus, child, and adult. Semin Pediatr Neurol 19:89–95.

52. Barton LL, Mets MB. 2001. Congenital lymphocytic choriomeningitis virus infection: decade of rediscovery. Clin Infect Dis 33:370–374.

53. Genbacev O, Larocque N, Ona K, Prakobphol A, Garrido-Gomez T, Kapidzic M, Bárcena A, Gormley M, Fisher SJ. 2016. Integrin α4-positive human trophoblast progenitors: functional characterization and transcriptional regulation. Hum Reprod 31:1300–1314.

54. Fisher S, Genbacev O, Maidji E, Pereira L. 2000. Human cytomegalovirus infection of placental cytotrophoblasts in vitro and in utero: implications for transmission and pathogenesis. J Virol 74:6808–6820.

55. Buchmeier MJ, Lewicki HA, Tomori O, Oldstone MB. 1981. Monoclonal antibodies to lymphocytic choriomeningitis and pichinde viruses: generation, characterization, and cross-reactivity with other arenaviruses. Virology 113:73–85.

56. Doench JG, Fusi N, Sullender M, Hegde M, Vaimberg EW, Donovan KF, Smith I, Tothova Z, Wilen C, Orchard R, Virgin HW, Listgarten J, Root DE. 2016. Optimized sgRNA design to maximize activity and minimize off-target effects of CRISPR-Cas9. Nat Biotechnol 34:184–191.

57. Ran FA, Hsu PD, Wright J, Agarwala V, Scott DA, Zhang F. 2013. Genome engineering using the CRISPR-Cas9 system. Nat Protoc 8:2281–2308.

58. Campeau E, Ruhl VE, Rodier F, Smith CL, Rahmberg BL, Fuss JO, Campisi J, Yaswen P, Cooper PK, Kaufman PD. 2009. A versatile viral system for expression and depletion of proteins in mammalian cells. PLoS One 4:e6529.

59. Sena-Esteves M, Tebbets JC, Steffens S, Crombleholme T, Flake AW. 2004. Optimized large-scale production of high titer lentivirus vector pseudotypes. J Virol Methods 122:131–139.

